# Entrainment of the *Arabidopsis* circadian clock to skeleton photoperiods

**DOI:** 10.64898/2026.10.21.760647

**Authors:** Stephanie S. I. Williams, Antony N. Dodd

## Abstract

Entrainment aligns the phase of the circadian clock and its outputs with 24 h fluctuations in the environmental conditions. One approach for investigating entrainment involves the application of skeleton photoperiods. In these protocols, light pulses are used to demarcate dawn and dusk, with the organism otherwise remaining under constant darkness. Historically, the consequences of skeleton photoperiod entrainment have been investigated by tracking behavioural responses in mammalian and insect models. In comparison, effects of skeleton photoperiod entrainment upon the plant circadian oscillator have received less attention. Using promoter-luciferase reporters for the *Arabidopsis thaliana* circadian clock, we monitored clock function during both entrainment to skeleton photoperiods, and subsequent free-running conditions. This enabled us to access molecular responses of the plant circadian oscillator to skeleton light conditions. We identified that individual circadian oscillator components in *Arabidopsis* can entrain to skeleton photoperiods. When these skeleton photoperiods were extended to represent long-day conditions, the oscillator underwent a phase jump relative to intact photoperiod entrainment such that it interpreted the dusk light pulse as dawn, and *vice versa*. This mirrors phase jumps that have been reported for rodent and insect behaviour following entrainment to long skeleton photoperiods, supporting the idea that phase jumps under skeleton photoperiod entrainment are a universal feature of circadian entrainment. Our results provide important context for the interpretation of physiological and developmental phenotypes of plants under modified light conditions.

## Introduction

Circadian clocks occur across the kingdoms of life, and have diverse evolutionary histories (Jabbur & Johnson, 2022). Their presence in such varied organisms reflects the selective advantage provided by circadian regulation (Ouyang et al., 1998; Woelfle et al., 2004; Dodd et al., 2005). To maintain an appropriate phase relationship between the clock, its outputs, and the external environment, circadian oscillators are continuously entrained to environmental cues known as zeitgebers. While the light/dark cycle is generally considered a reliable zeitgeber, mechanisms of light perception and transmission to the oscillator differ between organisms. Mammals possess a hierarchical timing system whereby molecular clocks located throughout the organism are coordinated by a central pacemaker in the suprachiasmatic nucleus (SCN) (Ko & Takahashi, 2006; Partch et al., 2014). The SCN clock receives light signals from the retina, with timing information communicated to peripheral clocks by endocrine cues and the autonomic nervous system (Dibner et al., 2010). In contrast, the plant circadian clock is semi cell-autonomous, so light perceived locally by a suite of photoreceptors directly regulates molecular clock components to achieve entrainment (Somers et al., 1998; Millar, 2003). In plants light also drives photosynthesis, which in the model plant *Arabidopsis thaliana* (Arabidopsis) entrains the clock through endogenous sugar signals (Haydon et al., 2013; Seki et al., 2017; Ohara et al., 2018). This dual usage of light by plants (for photoreceptor signalling, alongside photosynthesis) can complicate the interpretation of circadian phenotypes concerning plant responses to light conditions.

A consequence of entrainment to light is that the clock and associated pathways can also measure day length (photoperiod), enabling detection of and response to changes in the season. Plants are influenced strongly by the seasons, with photoperiod regulating diverse processes including the reproductive transition to flowering, seedling establishment and growth, primary metabolism, and bud dormancy (Bunning, 1960; Suárez-López et al., 2001; Niwa et al., 2009; Andrés & Coupland, 2012; Mengin et al., 2017; Singh et al., 2017).

Beyond plants, many insect species have photoperiodic responses. For example, in Drosophila ovarian diapause is regulated by photoperiod, with maturation occurring under long-day conditions (Saunders, 1997; Abrieux et al., 2020). Photoperiodism is also important for seasonal regulation of processes such as reproduction, migration, and hibernation in mammals, birds, and reptiles (Menaker & Keatts, 1968; Heldmaier et al., 1989; Gwinner, 1996; Rosa & Bryant, 2003; Williams et al., 2014). More recently, photoperiodic responses were identified in prokaryotes, with the cyanobacterial clock model *Synechococcus elongatus* PCC7942 having photoperiodic regulation of cold temperature tolerance (Jabbur et al., 2024).

One approach to probing the processes of entrainment and photoperiodism is to expose organisms to entrainment protocols known as “skeleton photoperiods.” In these protocols, dawn and dusk are defined by short periods of illumination, and the organism is otherwise maintained under conditions of continuous darkness. Mathematical modelling and biological experiments demonstrate that the circadian system can entrain successfully to these conditions, and that it exhibits bistability under skeleton photoperiod entrainment (Pittendrigh & Minis, 1964; Flôres et al., 2021). This bistability means that depending on the time elapsed between the skeleton light pulses, the clock can interpret either of the light pulses as dawn/dusk or dusk/dawn. In practice, this means that progressively altering the time elapsed between the two light pulses pushes the clock to a bifurcation, where the entrained phase jumps to a markedly different point within the 24 h cycle. This has been reported for insect eclosion (Pittendrigh & Minis, 1964; Saunders, 1979), rodent behaviour (Sharma et al., 1998; Evans et al., 2005; Comas et al., 2007; Spoelstra et al., 2014), flowering time in a short-day plant species (Oda, 1969), and an output of the clock in plants (Millar, 2003).

In plants, entrainment to skeleton photoperiods provides a strategy for overcoming their multiple and distinct uses of light, by balancing integrated photons across differing photoperiod lengths. However, information remains sparse about the entrainment of the Arabidopsis circadian oscillator skeleton photoperiods (Oda, 1969; Millar, 2003; Pokhilko et al., 2010; Bordage et al., 2016). Here, the experimental tools that are currently available in Arabidopsis have allowed us to directly examine the behaviour of individual clock components during skeleton photoperiod entrainment, adding depth to the previous examination of clock outputs that have been used in various organisms during skeleton photoperiod entrainment (Pittendrigh & Minis, 1964; Pittendrigh & Daan, 1976; Evans et al., 2005). We harnessed these tools to study the changes in entrainment that occur during exposure of plants to skeleton photoperiods, and how these conditions define the entrained phase of the circadian clock.

## Materials and Methods

### Plant material and growth conditions

*Arabidopsis thaliana* (Arabidopsis) genotypes used were Col-0, *CIRCADIAN CLOCK ASSOCIATED1::LUCIFERASE* (*CCA1::LUC*) (Pruneda-Paz et al., 2009), *GIGANTEA::LUCIFERASE* (*GI::LUC*) (J. Edwards et al., 2010), *TIMING OF CAB EXPRESSION1::LUCIFERASE* (*TOC1::LUC*) (Alabadí et al., 2001) and *DARK INDUCIBLE6::LUCIFERASE* (*DIN6::LUC*) (Frank et al., 2018). For cultivation, seeds were surface sterilised using 70% (v/v) ethanol for 10 minutes with agitation, washed 3 times with sterile distilled water, and resuspended in 0.1% (w/v) agar. Individual sterilised seeds were pipetted onto half strength Murashige and Skoog (MS) media with 0.8% (w/v) agar (pH 6.8), without added sucrose, then stratified in darkness at 4°C for before transfer into PHCBi MLR-352 plant growth chambers. Plants were grown under desired photoperiods at 19°C.

### Bioluminescence imaging of promoter-luciferase reporters

Seeds were prepared as above and stratified for 3 days, then grown on agar plates under short (8L:16D) or long-day (16L:8D) conditions for 11 days. On day 11 after stratification, plants were sprayed with 10 mM D-luciferin potassium salt solution (Biosynth) and returned to the same growth conditions.

At entrained dawn on the following day, seedlings were transferred to a bioluminescence imaging system comprising a Retiga LUMO camera (Teledyne Photometrics) mounted in a light-tight temperature-controlled incubator fitted with custom LED arrays for plant illumination. Both the camera and LED array were controlled using custom MicroManager 1.4 scripts. Plants were either maintained under the same intact short- or long-day light conditions, or transferred to corresponding short (3L:2D:3L:16D) or long-day skeleton photoperiod conditions (3L:10D:3L:8D) for 4 days (Figure 1). After monitoring luciferase reporters during this period of entrainment, free-running rhythms under constant light (LL) or constant dark (DD) were monitored for a further 3 days.

**Figure 1:**
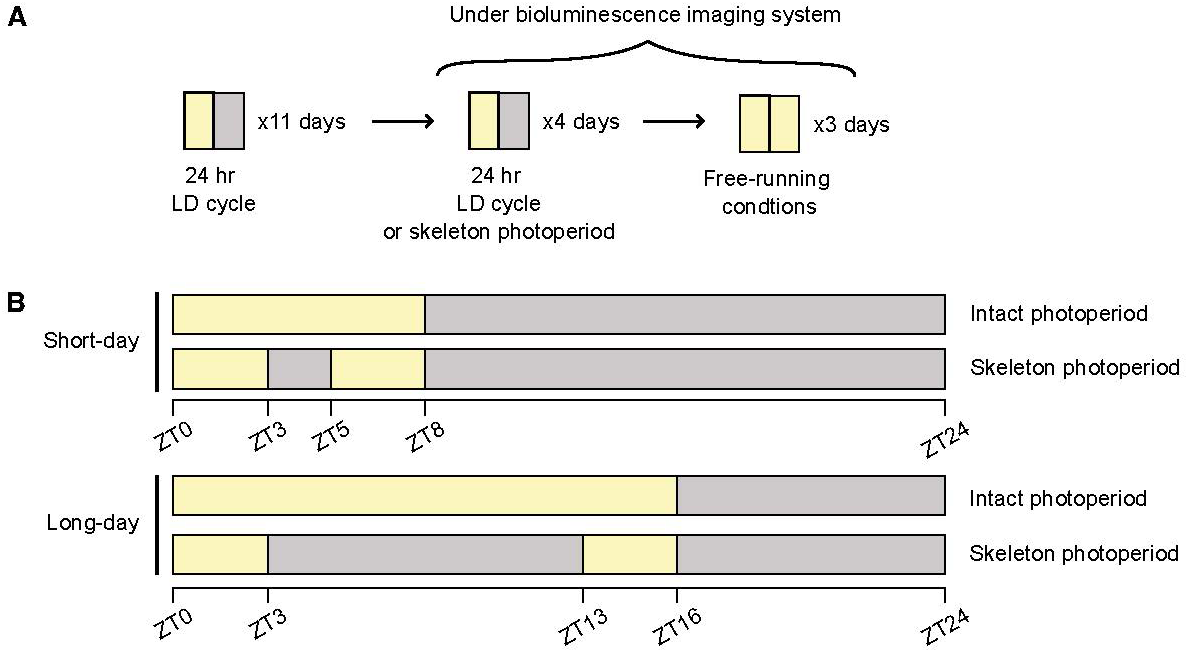
Experimental protocol used to explore entrainment characteristics of the Arabidopsis circadian clock to skeleton photoperiods. **(A)** Arabidopsis seedlings were initially entrained to short-day (8L:16D) or long-day (16L:8D) photoperiods for 11 days, before transfer to a bioluminescence imaging system. Using this, we monitored promoter activity of several core circadian oscillator components, alongside a clock output gene, for 4 days under light dark (LD) cycles. These cycles either matched the previous entrainment regime (intact photoperiod), or provided a corresponding skeleton photoperiod, as detailed in B. Following the entrainment cycles, plants were released into free-running conditions for 3 days.

All bioluminescence experiments began at entrained dawn (ZT0) and were performed at 19°C, with 20 _μm_ol m^-2^ s^-1^of red light and 20 _μm_ol m^-2^ s^-1^ of blue light (Figure S1). Bioluminescence images were captured at the end of each hour, using an exposure time of 3 minutes. Imaging data were processed using ImageJ (Schneider et al., 2012). To correct for camera noise, background signal was collected simultaneously and subtracted.

### Analysis of time-series data

To assess circadian parameters following entrainment, the free-running portion of each bioluminescence trace was analysed. The first 24 hours after transfer to constant conditions was excluded to remove the transient effects of the final dawn. To compare data across entrainment regimes and photoperiods, bioluminescence was normalised using z-scores.

Free-running data were analysed using eJTK and Fast Fourier Transform – Non-linear least squares method (FFT-NLLS), with no additional detrending in BioDare2 (Zielinski et al., 2014). Data collected under constant light were filtered for downstream analysis to include only seedlings that were statistically rhythmic (BH p < 0.01, eJTK), had a period between 20 and 28 hours (FFT-NLLS), and had ERR < 0.7. Phase was reported using circadian units, i.e., relative to the free-running period of the waveform (“modulo tau”). Circular means, angular deviation, rho, and Rayleigh’s p-value were calculated using the R package “circular” (Claudio Agostinelli & Lund, 2025). Comparisons of period estimates between intact and skeleton photoperiods was conducted using unpaired two-tailed Student’s t-tests. As circadian phase is a circular variable, standard linear statistics are not appropriate for its analysis. Therefore, to test whether phase differed between entrainment regime and photoperiod, each phase estimate was converted to radians and expressed as a pair of Cartesian coordinates (sine and cosine). This transformation allowed differences in phase means to be tested within a standard multivariate analysis of variance (MANOVA) model.

For each reporter, MANOVA models were fitted using *sin* and *cos* as bivariate response variables, and photoperiod, entrainment regime and their interaction as fixed factors. Pillai’s trace was used as the test statistic because it is robust to departures from multivariate normality and to heterogeneity of within-group variance. As the interactions were significant, entrainment regime was tested separately within each photoperiod (simple effects). This is in line with our *a priori* interest in the effect of entrainment regime on phase, rather than comparison between short and long photoperiods.

The bioluminescence peak height that occurred in response to dawn- and dusk-associated light pulses, under skeleton photoperiod entrainment, was defined as the maximum normalised bioluminescence that occurred within each window of illumination. Peak time was defined as the hour (0-24) within each 24 h imaging cycle that maximum bioluminescence was achieved.

### Hypocotyl elongation assays

To examine the effect of the entrainment regime on hypocotyl elongation, seeds were prepared as described previously and stratified for 5 days on vertically oriented ½ MS agar plates. Plates were then moved to LL for 48 h, to repress etiolation and induce germination. Following this, seedlings were grown under either short-day, long-day, short-day skeleton or long-day skeleton light conditions for 7 days. The next day, seedlings were photographed to enable measurement of hypocotyl length using ImageJ (Schneider et al., 2012). Hypocotyls were measured from the shoot–root junction to the shoot apical meristem (Belbin et al., 2019). To compare the hypocotyl length between groups, two-way ANOVA was used, followed by Tukey’s test for pairwise comparisons.

### Statistical analyses

Data are expressed as mean ± SEM unless otherwise stated. Sample sizes are indicated in figure legends, with each sample consisting of an individual seedling. All statistical analyses used R v4.4.1 (R Core Team, 2024). Analysis was run independently for each experimental replicate. Where appropriate, normality of residuals was checked using the Shapiro-Wilk test.

## Results

### Entrainment of the Arabidopsis circadian clock to skeleton photoperiods is daylength-dependent

To explore the dynamics of the plant circadian clock under skeleton photoperiods, we initially entrained *Arabidopsis* seedlings to either short (8L:16D) or long days (16L:8D) for 11 cycles, before transfer to a bioluminescence imaging system (Figure 1A). Using luciferase reporters, we monitored the promoter activity of several Arabidopsis circadian oscillator components including *CCA1* (Pruneda-Paz et al., 2009), *GI* (J. Edwards et al., 2010), and *TOC1* (Alabadí et al., 2001), alongside the clock output gene *DIN6* (Frank et al., 2018). The protocol involved monitoring luciferase activity for 4 days under light dark (LD) cycles that either (a) matched the previous entrainment condition (i.e. an intact photoperiod), or (b) comprised a skeleton photoperiod of equivalent daylength (Figure 1B). For skeleton photoperiods, we provided a 3-hour light pulse at dawn and a second 3-hour light pulse at dusk (Pokhilko et al., 2010). For plant models, skeleton light pulses must be considerably longer than for animal models to ensure that sufficient photosynthetic energy is provided for growth and survival. Following these entrainment cycles, plants were released into free-running conditions for a further 3 days. Initially, free-running conditions of constant light (LL) were used to prevent carbon starvation and maintain robust oscillations. Experiments were conducted twice independently; we present Experiment 1 in the main figures, focusing on information gained from the *CCA1::LUC* and *GI::LUC* reporters. Data for the *TOC1::LUC* and *DIN6::LUC* reporters, and Experiment 2, are in the supplemental figures.

Under intact short-(8L:16D) and long-day (16L:8D) LD cycles, both *CCA1::LUC* and *GI::LUC* had rhythmic bioluminescence, with *CCA1::LUC* activity peaking near dawn (Fig. 2A, B), and *GI::LUC* peaking near dusk (Fig. 2C, D). Consistent with previous reports, the shape of the oscillation differed slightly between the short- and long-day intact photoperiods (K. D. Edwards et al., 2010; Flis et al., 2016). Under LD cycles, activity of *TOC1::LUC* differed between short- and long-day intact photoperiods, with considerably lower amplitude oscillations under short-day conditions (Fig. S2A, B).

**Figure 2:**
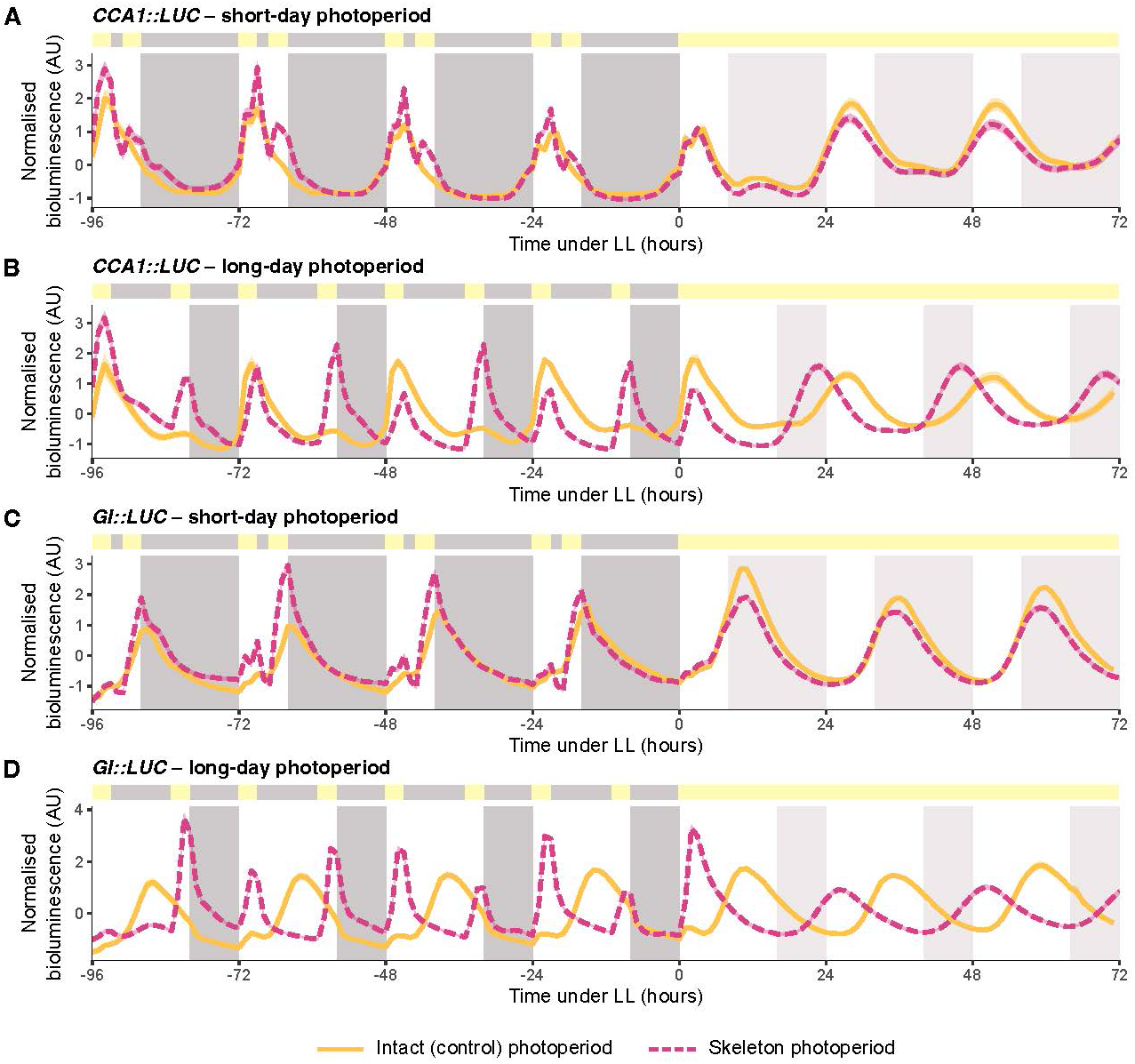
Entrainment of the *Arabidopsis* circadian clock to skeleton photoperiods is photoperiod dependent. Normalised bioluminescence of *CCA1::LUC* **(A, B)** and *GI::LUC* **(C, D)** seedlings under short-**(A, C)** or long-day **(B, D)** conditions, entrained to either intact (solid yellow) or skeleton (dashed pink) photoperiods for 4 cycles, then released into constant light (LL) at ZT0. For each panel, grey shading on the graph area indicates the light regime for intact photoperiods (darker grey represents night during entrainment; lighter grey represents subjective night under free-running conditions). The bar above each plot indicates the light regime of the skeleton photoperiod, with yellow representing lights on, and grey representing lights off. Data are mean (line) ± SEM (ribbon; *n* = 18-61 individual seedlings) and are from Experiment 1. Data from an independent, replicate experiment (Experiment 2) are in Figure S3.

Under short-day (3L:2D:3L:16D) and long-day skeleton photoperiods (3L:10D:3L:8D), *CCA1::LUC* and *GI::LUC* had two distinct peaks of expression during each 24-hour cycle; one in response to lights-on at dawn, and a second aligned to the second light pulse marking the end of the photoperiod (Fig. 2). This double-peaking has been reported previously for the response of *GI::LUC* to equinox skeleton photoperiods (Bordage et al., 2016). This also occurred for *TOC1::LUC* (Fig. S2) and was consistent with an independent, experimental replicate (Fig. S2 and Fig. S3).

Rhythms under the entraining LD cycles reflect the combined effects of the underlying endogenous circadian oscillator, and acute responses of clock reporters to light. We therefore also examined rhythms under conditions of constant light, as this enabled us to establish whether any after-effects were present once zeitgeber cycles were removed, as well as permit phase and period estimation. After entrainment to short-day conditions, the free-running phase under LL of *CCA1::LUC* (Fig. 3A), *GI::LUC* (Fig. 3B) and *TOC1:: LUC* (Fig. S4A) did not differ appreciably between intact and skeleton photoperiods (summarised in Table 1). In Experiment 1, under short days the phase was advanced slightly following skeleton entrainment compared with control conditions for *CCA1::LUC* (p = 0.02, -0.40 h; Fig. 3A), *GI::LUC* (p = 0.004, -0.39 h; Fig. 3B) and *TOC1::LUC* (p = 0.029, -0.48 h; Fig. S4A). In Experiment 2, phase was also slightly altered between intact and skeleton photoperiods for *CCA1::LUC* (p = 0.03, +1.35 h, Fig. S5A) and *TOC1::LUC* (p = 0.007, -0.84 h; Fig. S4B), but there was no effect of entrainment regime on *GI::LUC* phase (p = 0.964, Fig. S5B). The overall magnitude of these differences was very small, with angular deviation often overlapping between the two entrainment groups, suggesting broadly comparable phase distributions.

**Figure 3:**
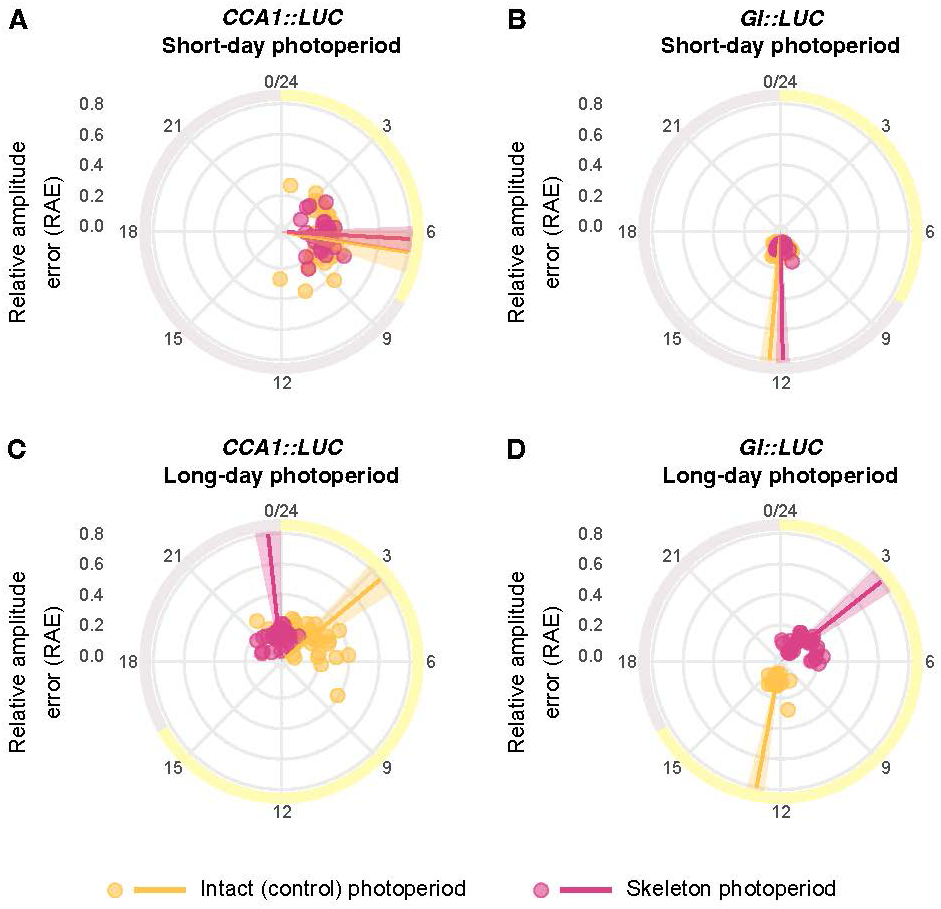
Phase is advanced by entrainment to long-day skeleton photoperiods. Circadian phase of *CCA1::LUC* **(A, C)** and *GI::LUC* **(B, D)** following entrainment to intact or skeleton photoperiods, determined using FFT-NLLS. Phase was calculated using the data collected between 24 - 72 h of free-running conditions, following 4 days of entrainment to either short-**(A, B)** or long-day **(C, D)** conditions under intact or skeleton photoperiods. Data are mean (line) ± angular deviation (ribbon; *n* = 18-61 individual seedlings represented as points) and are from Experiment 1. Yellow shading around each plot indicates subjective day, and grey shading indicates subjective night. The radial axis indicates the relative amplitude error (RAE). Data from an independent replicate experiment (Experiment 2) are in Figure S5.

**Table 1:**
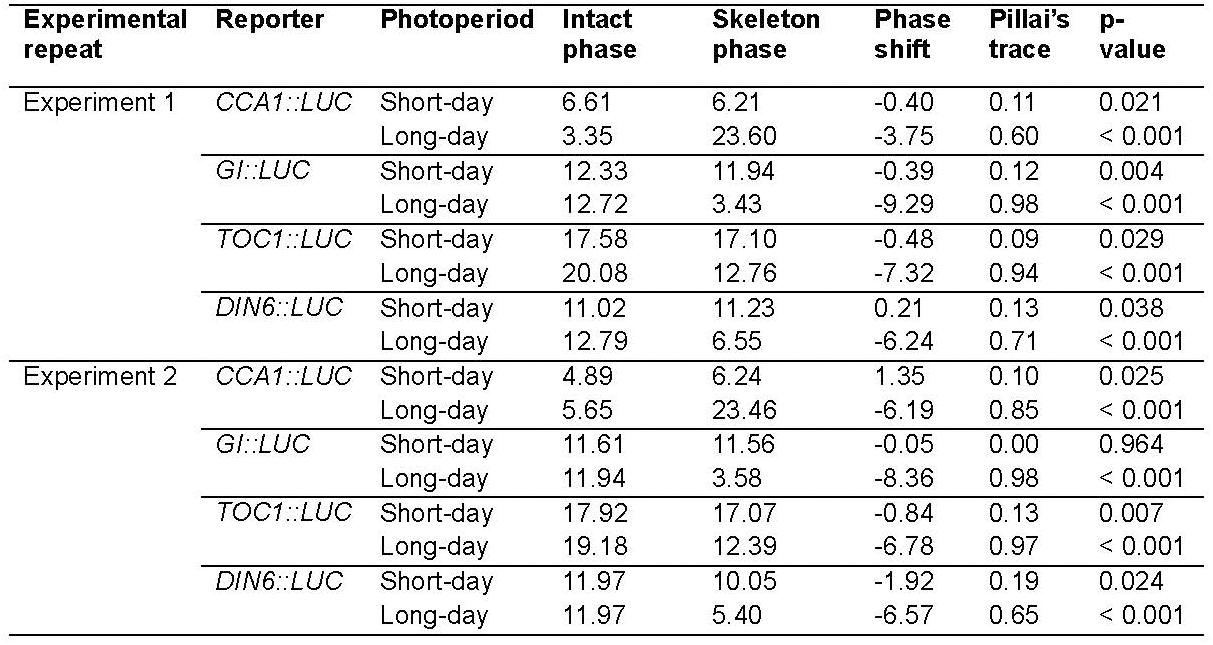
Phase shifts caused by entrainment of Arabidopsis to skeleton photoperiods. Mean circadian phase per entrainment regime determined with FFT-NLLS. “Phase shift” is the phase difference between intact and skeleton photoperiod entrainment, where negative values indicate a phase advance. Statistical comparisons of phase performed using MANOVA with Pillai’s trace as the test statistic (higher values indicate a stronger multivariate effect of entrainment regime; range 0-1). The associated p-values indicate whether phase differs significantly between intact and skeleton entrainment regimes. Data from two independent replicate experiments.

In contrast, after entrainment to long-day conditions, the free-running phase under LL differed substantially between intact photoperiods and skeleton photoperiods. This occurred for *CCA1::LUC* (p < 0.001, averaging -5 h across experiments; Fig. 3C), *GI::LUC* (p < 0.001, averaging -8.8 h across experiments; Fig. 3D) and *TOC1::LUC* (p < 0.001, averaging -7.1 h across experiments, Fig. S4A, B). In each case, the phase was advanced in response to skeleton entrainment. The direction and magnitude of this phase shift was comparable across *CCA1*, *GI*, and *TOC1,* suggesting a systematic change in oscillator phase, rather than a gene-specific effect (Table 1).

To rule out that this phase divergence being driven by altered period and to check for stable entrainment, we evaluated the free-running period. Across reporters, the entrainment regime had no major effect on free-running period with minor exceptions (Fig. S6A, B). For instance, the period of *TOC1::LUC* was significantly longer following entrainment to long-day skeleton compared with the control in both experimental replicates (Fig. S6A, B). These differences are sufficiently modest (up to 40 minutes) meaning that they are unlikely to be of physiological relevance. Instead, this may reflect the sensitivity of period estimation and our relatively large sample size (Table S1). Overall, period estimates across all groups did not markedly differ from 24 hours, suggesting the circadian clock entrained stably to intact and skeleton photoperiod regimes across both long and short days and that phase differences were not due to changes in period.

Considered together, these experiments support the idea that entrainment to short-day skeleton photoperiods preserves the entrained and free-running dynamics of intact short-day photoperiods. In contrast, entrainment of Arabidopsis to long-day skeleton photoperiods substantially shifted the circadian phase compared with intact photoperiod conditions.

### Entrainment to long-day skeleton photoperiods switches the circadian phase between the two light pulses

The divergence in free-running phase under long-day, but not short-day, conditions raised the question of whether this shift occurred abruptly or developed progressively over successive entrainment cycles (Dixon et al., 2014). To address this, we quantified the peak bioluminescence associated with the “dawn” and “dusk” skeleton light pulses (peak 1 and peak 2, respectively) in each 24 h cycle. Although these pulses partly mask the underlying circadian rhythm, they fall at distinct circadian phases and are identical in intensity and quality across all the entraining cycles, so we attribute any change in their relative height to a change in clock state. Under short-day skeleton photoperiods, the relative height of peak 1 and peak 2 remained stable across all four entrainment cycles. For *CCA1::LUC*, peak 1 was always greater than peak 2 (Fig. 4A) and for *GI::LUC*, peak 2 was consistently greater than peak 1 (Fig. 4B). Under long-day skeleton photoperiods, the relative height of the two peaks changed progressively over time, so that by the end of the four cycles of entrainment, the dominant peak switched between the two light pulses. The timing of this switch differed between the clock reporters. For *CCA1::LUC*, the switch from the dominance of peak 1 to peak 2 occurred between cycle 1 and cycle 2 (i.e. within 48 h of the experiment start, Fig. 4C). The switch of *GI::LUC* dominance from peak 2 to peak 1 occurred between cycles 2 and 3 (within 72 h of the experiment start, Fig. 4D). These transitional dynamics also occurred in a second independent experiment (Fig. S7).

**Figure 4:**
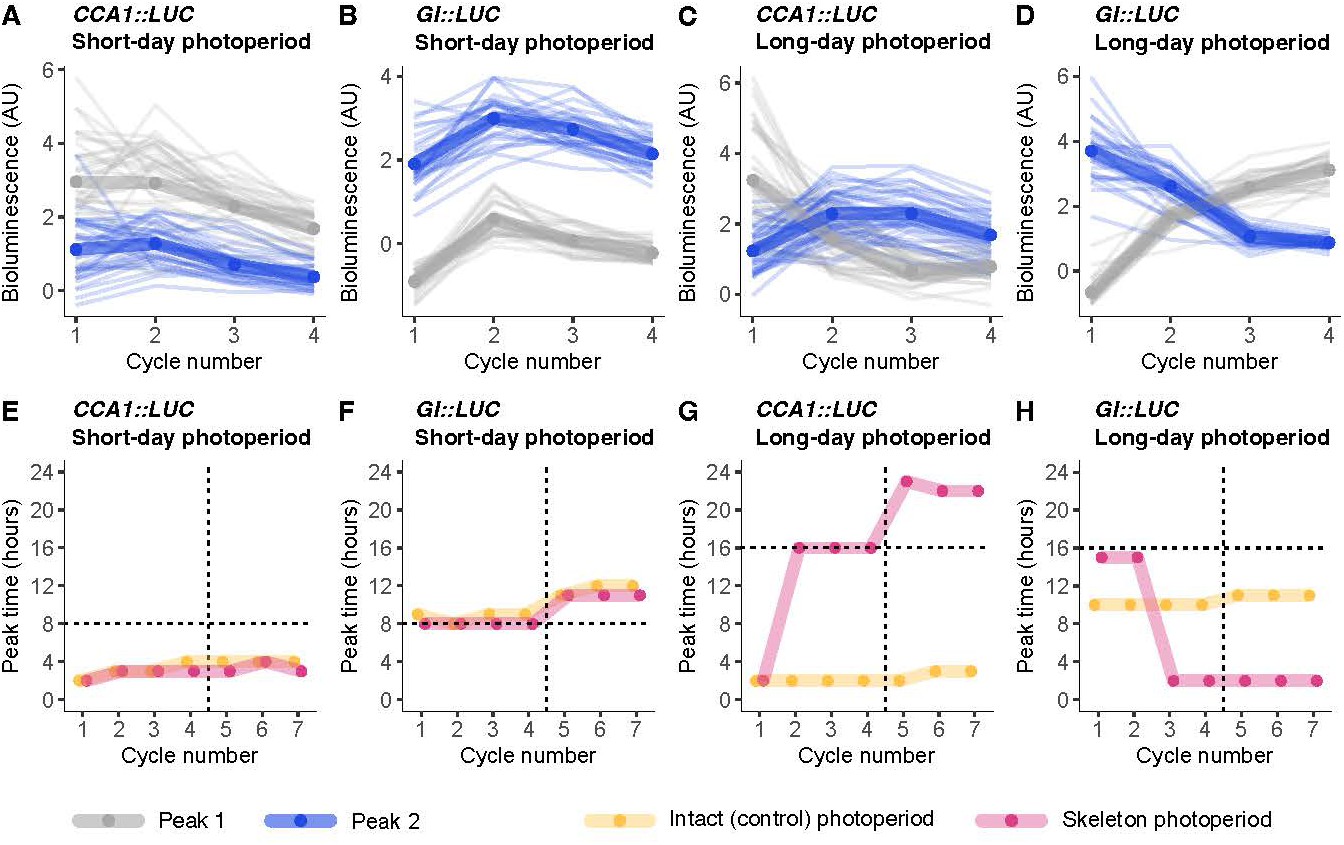
Entrainment to long-day skeleton photoperiods switches the circadian phase between the two light pulses. (A-D) Peak bioluminescence measurements of *CCA1::LUC* and *GI::LUC* associated with dawn (peak 1; grey) dusk (peak 2; blue) light pulses, measured under both short-and long-day skeleton photoperiods. Each line represents an individual seedling (*n* = 18-61), with the mean overlayed on top. **(E-H)** Examination of how these dynamics are reflected during entrainment and free-run under both intact and skeleton photoperiods. Data show timing of maximum daily bioluminescence of *CCA1::LUC* and *GI::LUC* across each 24 h window during both entrainment and subsequent free-run in constant light. Daily maxima may be due to either peak 1 or peak 2. Vertical dashed lines indicate the transition from entrainment to free-run: cycles 1-4 were the entrainment period, and cycles 5-7 were free-running conditions of constant light. Horizontal dashed lines indicate the end of the light period for intact photoperiods, and end of the second light pulse for skeleton photoperiods. Peak time data are mean of replicate seedlings. Data derived from Experiment 1. Equivalent graphs from an independent replicate experiment (Experiment 2) provided as Figure S7.

To quantify the extent to which these dynamics represent changes in the phase, we tracked the timing of maximum bioluminescence activity within each 24 h cycle, during both entrainment and LL. Under short-day entrainment, the timing of the peak of *CCA1::LUC* and *GI::LUC* was unaltered between plants entrained to intact and skeleton photoperiods (Fig. 4E, F respectively). By comparison, under long-day conditions, we observed a shift in the timing of peak activity of *CCA1::LUC* and *GI::LUC* during entrainment to skeleton photoperiods relative to intact photoperiods. On the second day of long-day LD cycles, the *CCA1::LUC* maxima switched from being the dawn-timed peak occurring 3 h after dawn, to the dusk-timed peak (16 h after dawn; Fig. 4G). The reverse was true for *TOC1::LUC* (Fig. S8A, B). This peak timing switch also occurred for *GI::LUC* from the third day of LD cycles, where the maxima jumped from 15 h after dawn (aligned to the dusk-timed light pulse) to 2 h after dawn (aligned to the dawn-timed light pulse; Fig. 4H). This also occurred in our independent replicate experiment (Fig. S7 and Fig. S8), demonstrating that just a few cycles of long-day skeleton photoperiods are sufficient to cause a phase jump within the Arabidopsis circadian clock.

### Free-run in constant darkness supports the differing phase of entrainment under intact and skeleton photoperiods

Given that light is an important zeitgeber for the Arabidopsis circadian clock (McClung, 2006), we hypothesised that acute responses to light at dawn, upon release into free-running conditions, may obscure the phase established by the skeleton photoperiods. Therefore, we repeated our experimental design but, after the 4 days of LD cycles, released plants into constant darkness (DD) instead of constant light (LL).

Reporters of the Arabidopsis circadian clock damp rapidly under DD, so it is possible to study their behaviour for only a few days (Dalchau et al., 2011). *CCA1::LUC* activity decreased rapidly under DD (Wu et al., 2008) and had no clear rhythmicity, so it was not possible to extract accurate phase estimates (Fig. 5A-D). *GI::LUC* was rhythmic under DD, albeit with lower amplitude and peak bioluminescence than during entrainment (Fig. 5E-H).

**Figure 5:**
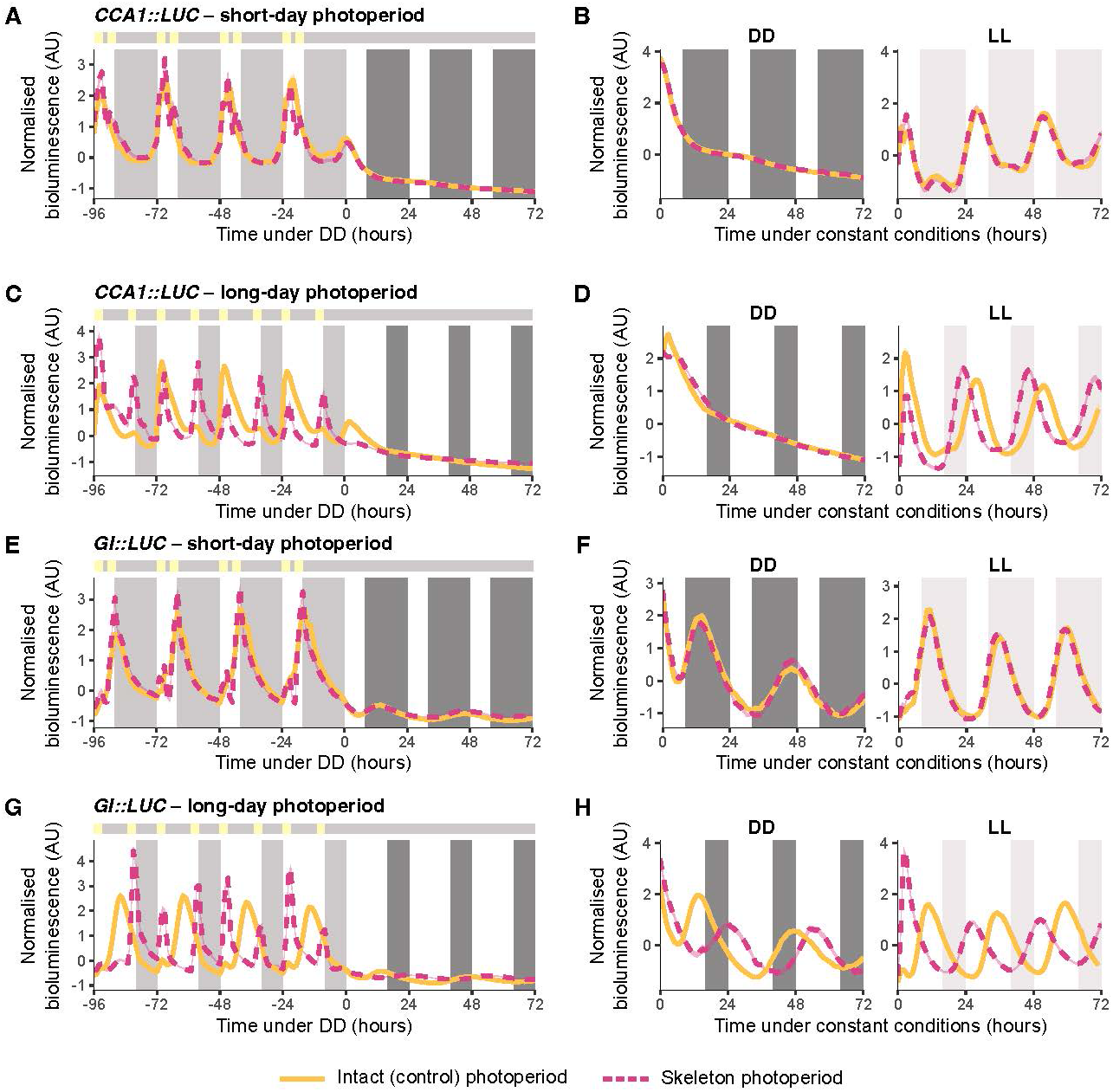
The differing phases of clock reporters under long-day intact and skeleton photoperiods is confirmed under free-running conditions of constant darkness. Normalised bioluminescence of *CCA1::LUC* (A-D) and *GI::LUC* (E-H) seedlings under short-**(A, B, E, F)** or long-day **(C, D, G, H)** conditions, using either intact (solid yellow) or skeleton (dashed pink) photoperiods. Following entrainment, plants were released into free-running conditions of constant darkness (DD) at ZT0. For panels **A**, **C**, **E,** and **G**, shading on the plot area indicates the light regime of the intact photoperiod, with light grey bars night during entrainment, and dark grey indicating subjective night under DD. The bar above each plot indicates the skeleton photoperiod regime, with yellow indicating lights on and grey indicating lights off. **(B**, **D**, **F**, **H)** Re-normalised data from the DD portion of **A, C, E, G**. Under DD *CCA1::LUC* in panels **B** and **D** was not rhythmic. LL sections of these panels are re-normalised data from Figure 2, for comparison purposes. Data are mean (line) ± SEM (ribbon; *n* = 24-39 individual seedlings) from Experiment 1. Data from an independent replicate experiment (Experiment 2) in Figure S10.

To examine rhythmicity and peak timing more clearly, we isolated the DD free-running portion of the signal and normalised these data independently of the LD cycles.

Under DD, rhythms of *GI::LUC* activity were largely unaltered by prior entrainment to short-day skeleton photoperiods, compared with intact photoperiods (Fig. 5E, F). In contrast, under long-day conditions, there was a clear phase shift caused by skeleton photoperiod entrainment (Fig. 5G, H; Fig. S9). Specifically, after entrainment to intact long-day photoperiods, the first peak in *GI::LUC* under DD occurred 13 h after subjective dawn (37 h after the last dawn during entrainment, Fig. 5G, H). In comparison, after long-day skeleton photoperiod entrainment, the first peak occurred 24 h after subjective dawn (35 h after the light onset of the final dusk skeleton pulse; Fig. 5G, H). This occurred also in a second independent experiment (Fig. S10). The time to peak bioluminescence under DD was more similar between long intact and skeleton photoperiods, when accounting for the final light onset. This suggests that for long skeleton photoperiods, the dusk-associated light pulse has the greater influence on the entrained phase of the clock.

Similar to LL, the free-running period of *GI::LUC* under DD was unaltered by skeleton photoperiod entrainment (Fig. S9) indicating that the phase change was not due to a difference in period length. The free-running period was considerably longer under DD (closer to 32 h, Fig. S9A) than LL (∼24 h), which is consistent with previous reports and Aschoff’s rule (Aschoff, 1960; Millar et al., 1995; Somers et al., 1998).

This combination of phase shift and swapped alignment of the peak time to each light pulse strongly supports the notion of a phase jump in plants entrained to long-day skeleton photoperiods, with the 10-hour dark period between the two light pulses – once interpreted as daytime – now being interpreted as night by the clock. In contrast, under short-day conditions, *GI::LUC* activity under DD did not differ between skeleton- and control-entrained plants, indicating that no such phase jump occurred.

### DIN6::LUC reports skeleton photoperiod entrainment downstream of the circadian oscillator

We also investigated the promoter activity of *DIN6*, which is a circadian clock-regulated marker for sugar signalling thought to be positioned downstream of the oscillator (Frank et al., 2018). Across all skeleton and control photoperiod conditions, the dynamics of *DIN6::LUC* were more variable compared with the circadian oscillator reporters. This was indicated by a greater spread of free-running period (Figure S6), phase, and angular variance (Figure S4C, D). Following entrainment to short-day conditions, there was a small difference in circadian phase between intact and skeleton photoperiod entrainment, with broadly comparable phase distributions (p < 0.05; Figure S5). Similar to reporters for circadian clock components, *DIN6::LUC* phased significantly earlier following entrainment to long-day skeleton photoperiods compared with control (intact) photoperiods (p < 0.001, averaging -6.4 h phase difference across experiments, Table 1). In comparison to clock reporters, we did not observe a change in the peak time of promoter activity for *DIN6::LUC* between long intact and skeleton photoperiods (Fig. S7), supporting the notion that the *DIN6* promoter integrates circadian signals with other metabolic signals (Frank et al. 2018).

However, the shift in entrained phase of *DIN6::LUC* demonstrates that there was a skeleton photoperiod-induced phase jump for this output of the clock, as well as for clock components.

## Discussion

### Skeleton photoperiods can produce jumps in the entrained phase of Arabidopsis plants

Here, we identified that skeleton photoperiods can recapitulate short-day, but not long-day, entrainment of the Arabidopsis circadian clock, measured using several reporters of circadian oscillator components. This phase change between long-day skeleton versus intact photoperiods was predicted by Pittendrigh (1981), and has been experimentally-demonstrated by monitoring circadian-regulated behaviour of organisms including rodents and insects (Pittendrigh & Minis, 1964; Pittendrigh & Daan, 1976; Evans et al., 2005).

However, this feature of entrainment has received comparatively little attention in plants. Our data provide experimental evidence from Arabidopsis to support the occurrence of the phase jump, and this concept of bistability in the entrainment of circadian oscillators. Moreover, they show at a high temporal resolution the direct impact of these entrainment regimes on clock oscillator components.

By using tools available in Arabidopsis (Millar et al., 1992), we detected this phase jump under long-day skeleton photoperiods through direct monitoring of the circadian oscillator components, rather than through behavioural (e.g. Pittendrigh & Minis, 1964; Pittendrigh & Daan, 1976; Evans et al., 2005), or genetic outputs of the clock (Millar, 2003). *CCA1*, *GI,* and *TOC1* represent temporally-distinct components of the Arabidopsis circadian oscillator and form parts of its morning and evening loop regulation (Hsu & Harmer, 2014). Each of these reporters differs in its sensitivity to photoperiod and light quality – for example, *CCA1* is also regulated strongly by light signalling. Each of these genes undergoes comparable shifts in phase (both direction and magnitude), demonstrating a systematic change in the circadian clock, rather a gene-specific, oscillator loop-specific, or light sensitivity-effect. We observed that under long days, following release into LL, all reporters tested showed advanced phase (Figure 2, 3) caused by skeleton entrainment, whilst under DD (for *GI*) there was a phase delay (Figure 5). This reveals that light at dawn, upon release into LL, influences the phase of skeleton-entrained plants. Assuming that our data extend to other plants, our results likely explain why the photoperiodic control of flowering in *Lemna perpusilla* (duckweed) is disrupted by entrainment to skeleton photoperiods (Oda, 1969).

Continuous monitoring of the oscillator during our experiments allowed investigation of changes in clock regulation under both entrainment and free-running conditions. During skeleton photoperiod entrainment, all reporters tested had two distinct peaks during each 24 h cycle, with these coinciding with the time of each light pulse. Further examination of the peaks or *CCA1* and *GI* under long days revealed that at the beginning of the experiment (Cycle 1, Figure 5C, D), one of these peaks dominated: either the first peak (for *CCA1::LUC*), or the second peak (for *GI::LUC*). However, with each successive 24 h cycle of skeleton entrainment, the primary peak diminished and the secondary peak became increasingly dominant. We reason that there could be several explanations for this. Our bioluminescence imaging method for monitoring the clock measures population of cells across multiple tissues within each seedling. Given that in plants, the circadian oscillator is semi cell-autonomous (Nakamichi et al., 2003; Yakir et al., 2011; Wu & Locke, 2026), it is possible that there was some stochasticity in the entrainment of the clock of individual cells, such that phase jumps occurred in different cells during different 24 h cycles. This could also be occurring at level of individual tissues, given that differing plant tissue types have differing circadian dynamics (Thain et al., 2002; Endo et al., 2014; Greenwood et al., 2019). Taken as a whole plant average (Paajanen et al., 2021), this would produce a progressive change over time in the dominant peak within the clock cycle. In either case, this could suggest that individual cells or cell types have somewhat variable entrainment characteristics. A further explanation is that the clock gates an acute response to light of each of these reporters, and a change in entrainment of the underlying oscillator shifts the window of time that the reporters have greatest light responsiveness. Such acute light responses tend to occur during entrainment of plants to square-wave light/dark cycles but are less apparent under field conditions (Nagano et al., 2019; Dantas et al., 2021). A further explanation is that the complexity of the interlocking feedback loops of the Arabidopsis circadian oscillator permits a degree of flexibility under entrainment conditions. Therefore, the shifts in peak height could be a consequence of the differing transient effects that arise between components whilst the oscillator shifts from one stable condition to another (Dixon et al., 2014). A previous study using skeleton entrainment to equinox photoperiods found no progressive change over time in light-induced peak height of an oscillator component (Bordage et al. (2016)), further supporting our finding that the change in peak dominance occurs only under conditions that elicit a jump in the entrained phase. Future experimental work could quantify the number of cycles of skeleton photoperiod entrainment that is required to achieve a new steady-state within the circadian oscillator.

### Implications for plant physiology and development

Physiological and developmental investigations in plants sometimes make use of skeleton photoperiods (e.g. Oda, 1969; Graf et al., 2010), while others studies have used conceptually similar light pulse protocols called “night break” experiments to manipulate photoperiod conditions (Pearce et al., 2017). One advantage of these approaches is that they enable manipulations with different photoperiod lengths to deliver the same total amount of integrated photosynthetic light over each 24 h cycle. Our data suggest that under conditions where long skeleton photoperiods are used, physiological or developmental phenotypes might differ from those that are expected because of phase jumps in the circadian oscillator. If such phenotypes arise, it is important to monitor the circadian clock to determine whether the altered phenotype could be associated with phase jumping.

We tested this possibility by comparing hypocotyl elongation under skeleton and intact photoperiods. The hypocotyl is the stem of the germinating seedling, and it normally undergoes enhanced elongation under short photoperiods compared with long photoperiods. (Seaton et al., 2015; Martín et al., 2018). There was no difference in hypocotyl length between short-day intact and skeleton photoperiods (Fig. S11A), whereas long-day skeleton photoperiods appeared to induce a phenotype known as etiolation (often identifiable by chlorotic colouring and exaggerated elongation (Jedynak et al., 2022)) (Fig. S11B). While both short and long-day skeleton photoperiod conditions delivered the same total duration of light, only long-day skeletons produced a phenotype associated with light deprivation. Therefore, a combination of proper circadian entrainment and light duration is required to elicit certain photoperiod-regulated behaviours.

A further experimental context could relate to the examination of circadian period length mutants of Arabidopsis under long skeleton photoperiods. In this case, our interpretation is that long period mutants would require a longer duration between the first and second skeleton light pulses to elicit a phase jump, compared with the wild type.

### Considerations for entrainment under natural and artificial light conditions

Our results have implications for how plants interpret complex or fluctuating light conditions in the field. Under natural conditions, situations may arise whereby plants become exposed to light during the dark period, in a manner reminiscent of skeleton photoperiod conditions. One example of this is transient exposure to artificial light sources such as street illumination and satellite-induced skyglow (Kocifaj et al., 2021; Gaston & de Miguel, 2022). Our data suggest that if such light is of sufficient intensity and duration to act upon entrainment (1-5 _μm_ol m s light can alter the Arabidopsis free running period (Somers et al., 1998)), changes in entrained phase might be induced. This could have implications for plant physiology and phenology and may be worthy of future investigation to understand how artificial light sources might impact plant ecology and agriculture.

## Author contributions

SW conducted experimentation, data analysis, and data visualization, SW and AD interpreted the data, and SW and AD wrote the paper.

## Conflicts of interest

The authors declare no potential conflicts of interest with respect to the research, authorship, and/or publication of this article.

## Acknowledgements

We thank Andrew Millar, Maria Luísa Jabbur, and Luíza Lane de Barros Dantas for helpful feedback and suggestions; Maximillian Jones for feedback on the manuscript; and Phil Robinson for help with photography.

## Funding

This work was funded by UKRI-BBSRC (Institute Strategic Programme BRiC BB/X01102X/1), the John Innes Centre, the John Innes Foundation, and the European Union (ERC, Synergy award 101166968 “MicroClock”); views and opinions expressed are those of the authors only and do not necessarily reflect those of the European Union or the European Research Council Executive Agency.

## Figure legends

## Notes

### Competing Interest Statement

The authors have declared no competing interest.

