## Supplementary Information for "Entrainment of the *Arabidopsis* circadian clock to skeleton photoperiods"

**Supplementary figures:**

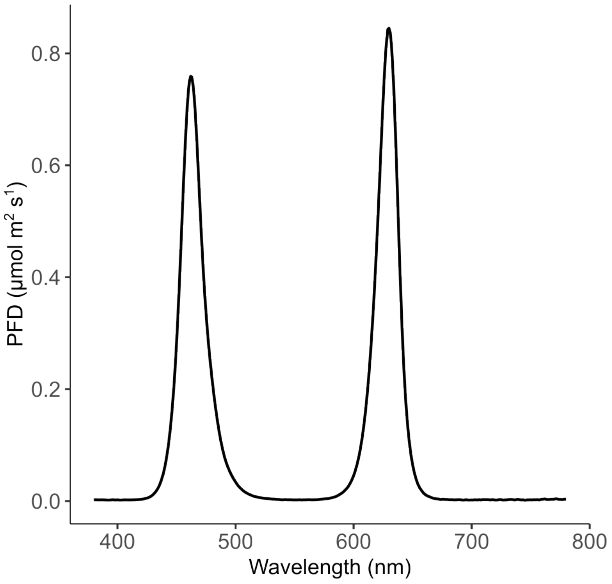


**Figure S1: Spectral distribution of LED panel used during bioluminescence imaging experiments.** Measured as photon flux density (PFD).

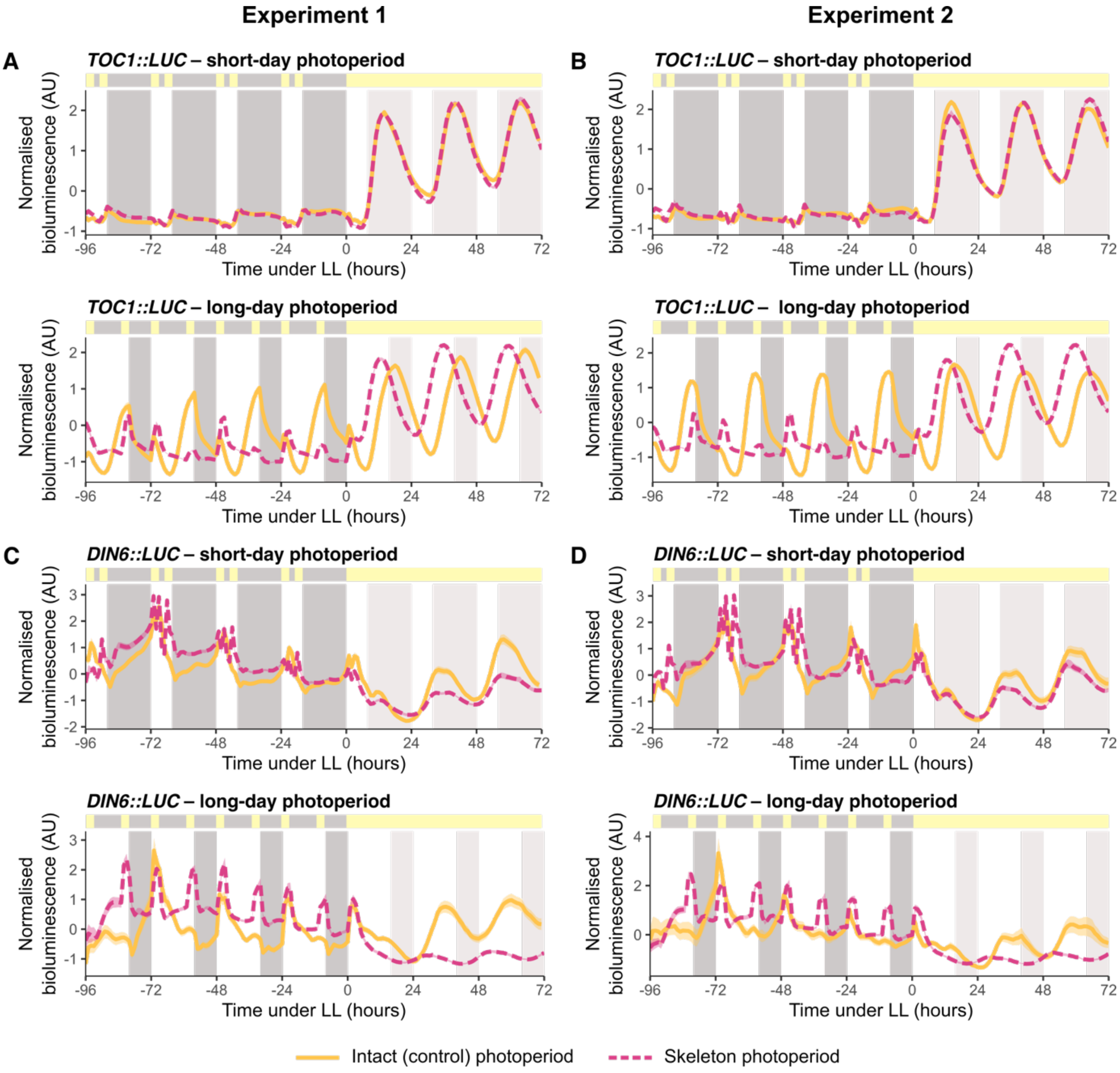


**Figure S2: Further bioluminescent reporters demonstrate entrainment of the *Arabidopsis* circadian clock to skeleton photoperiods is photoperiod dependent.** Normalised bioluminescence of circadian clock component *TOC1::LUC* **(A, B)** and downstream clock output *DIN6::LUC* **(C, D)** seedlings under short- or long-day conditions, entrained to either intact (solid yellow) or skeleton (dashed pink) photoperiods for 4 cycles, and released into constant light (LL) at ZT0. For each panel, grey shading on the graph area indicates the light regime for intact photoperiods (darker grey indicates night during entrainment; lighter grey indicates subjective night under free-running conditions). The bar above each plot indicates the light regime of the skeleton photoperiod, with yellow representing lights on and grey representing lights off. Data are mean (line) ± SEM (ribbon; *n* = 10-41 individual seedlings), from two independent experiments. **(A, C)** Experiment 1; **(B, D)** Experiment 2.

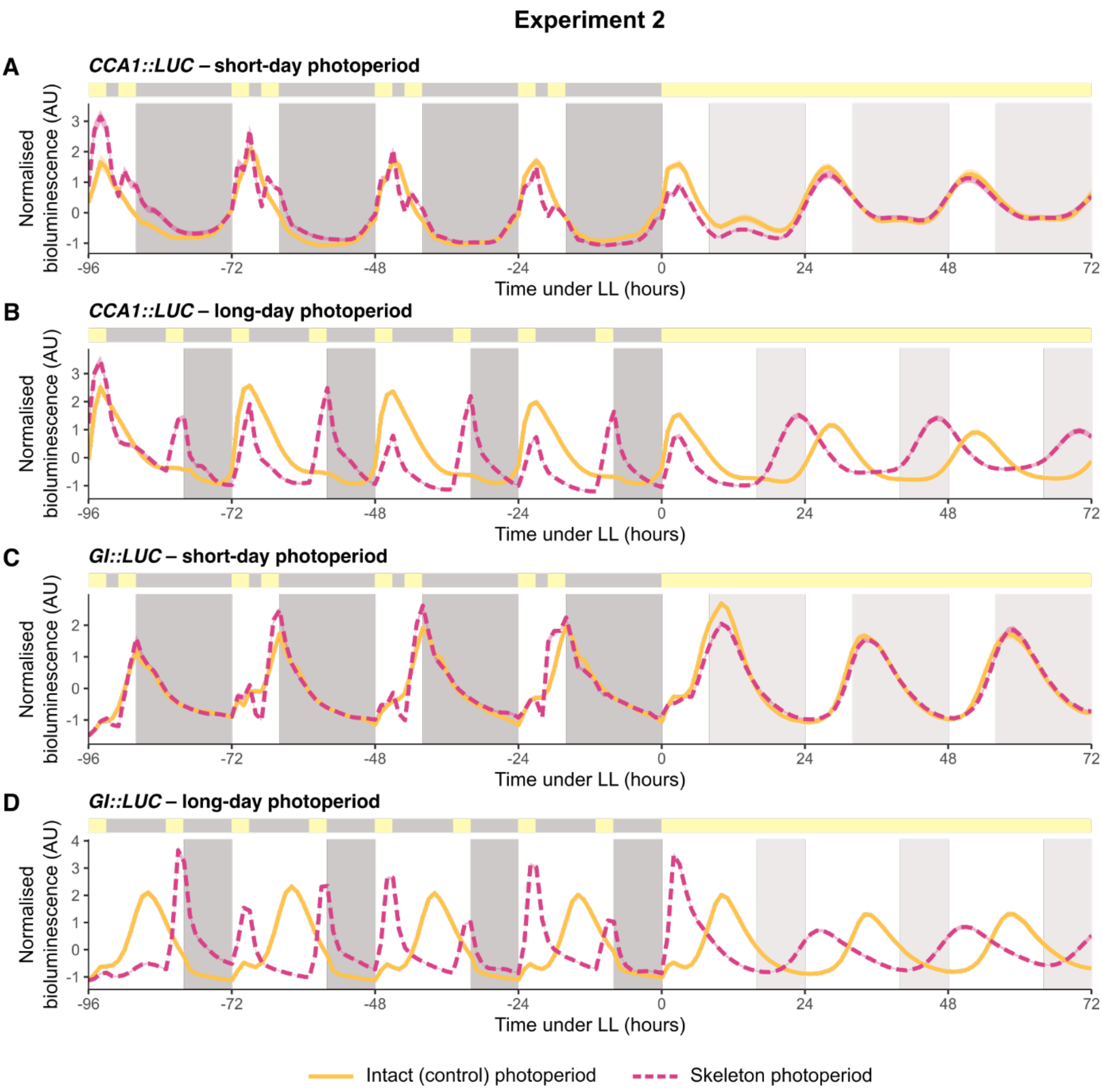


**Figure S3: Entrainment of the *Arabidopsis* circadian clock to skeleton photoperiods is photoperiod dependent.** Normalised bioluminescence of **(A, B)** *CCA1::LUC* and **(C, D)** *GI::LUC* seedlings under **(A, C)** short- or **(B, D)** long-day conditions, entrained to either intact (solid yellow) or skeleton (dashed pink) photoperiods for 4 cycles, and released into constant light (LL) at ZT0. For each panel, grey shading on the graph area indicates the light regime of the intact photoperiod (darker grey indicates night during entrainment; lighter grey indicates subjective night under free-running conditions). The bar above each plot indicates the light regime of the skeleton photoperiod (yellow indicates lights on; grey indicates lights off). Data are mean (line) ± SEM (ribbon; *n* = 17-41 individual seedlings) and derive from an independent, replicate experiment (Experiment 2).

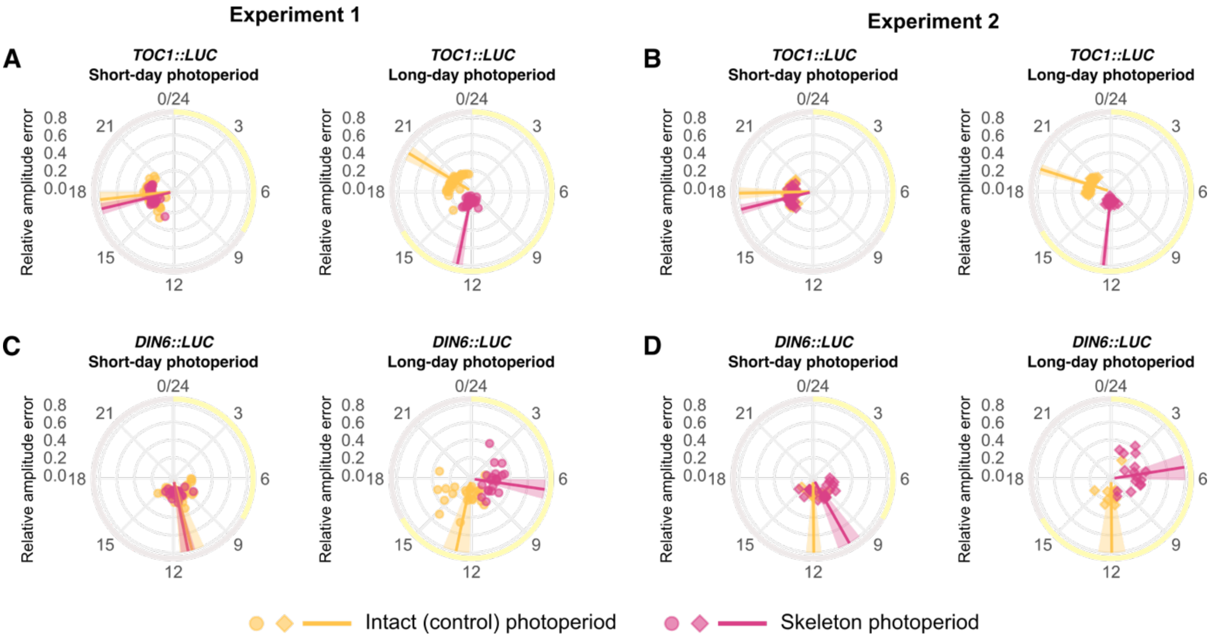


**Figure S4: Further bioluminescent reporters demonstrate phase advancement by entrainment to long-day skeleton photoperiods.** Circadian phase of circadian clock component *TOC1::LUC* **(A, B)** and downstream clock output *DIN6::LUC* **(C, D)** following entrainment to intact or skeleton photoperiods determined using FFT-NLLS. Phase was calculated using data collected between 24 - 72 h free-running conditions, following 4 days of entrainment to either short- or long-day conditions under intact or skeleton photoperiods. Data are mean (line) ± angular deviation (ribbon; *n* = 10-41 individual seedlings represented as points) and derive from Experiment 1 **(A, C)** and Experiment 2 **(B, D)** (an independent repeat). Yellow shading around each plot indicates subjective day, and grey shading indicates subjective night for intact photoperiod entrainment. Radial axes indicate the relative amplitude error (RAE).

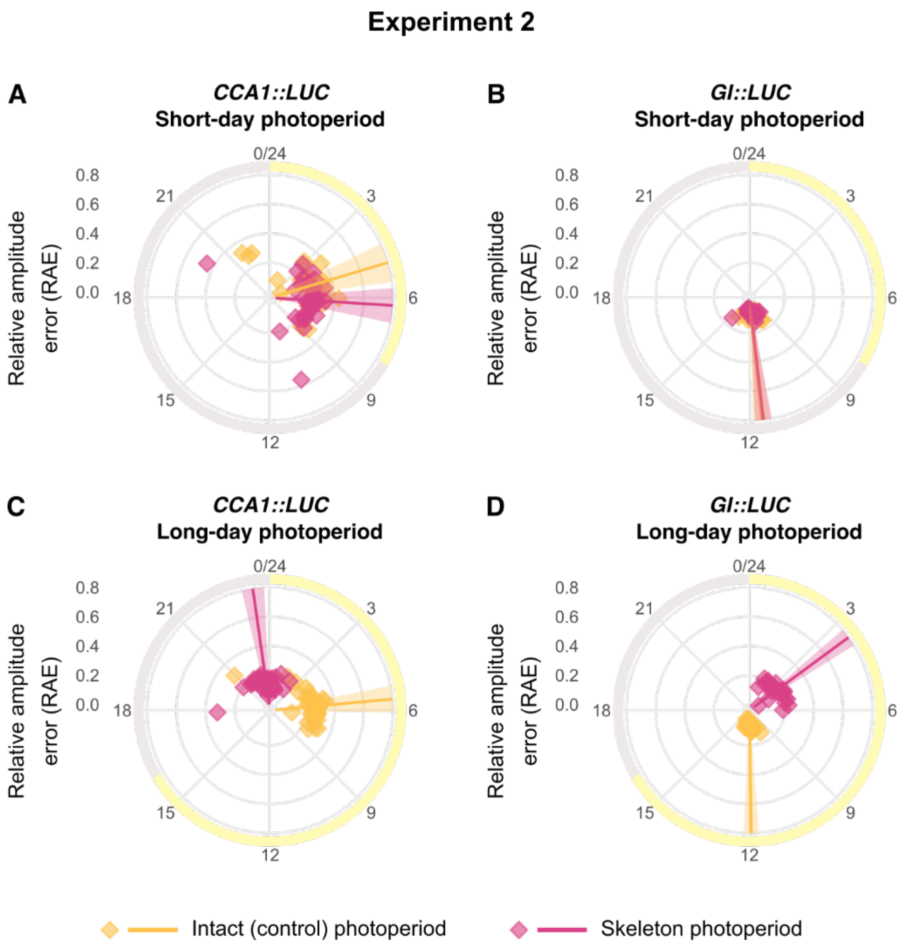


**Figure S5: Phase is advanced by entrainment to long-day skeleton photoperiods.** Circadian phase of *CCA1::LUC* **(A, C)** and *GI::LUC* **(B, D)** following entrainment to intact or skeleton photoperiods, determined using FFT-NLLS. Phase was calculated using data collected between 24 - 72 h of free-running conditions, following 4 days of entrainment to either short- **(A, B)** or long-day **(C, D)** conditions, under intact or skeleton photoperiods. Data are mean (line) ± angular deviation (ribbon; $n$ = 17-41 individual seedlings represented as points) and are from Experiment 1. Yellow shading around each plot indicates subjective day, and grey shading indicates subjective night. The radial axis indicates the relative amplitude error (RAE). Data are from an independent replicate experiment (Experiment 2).

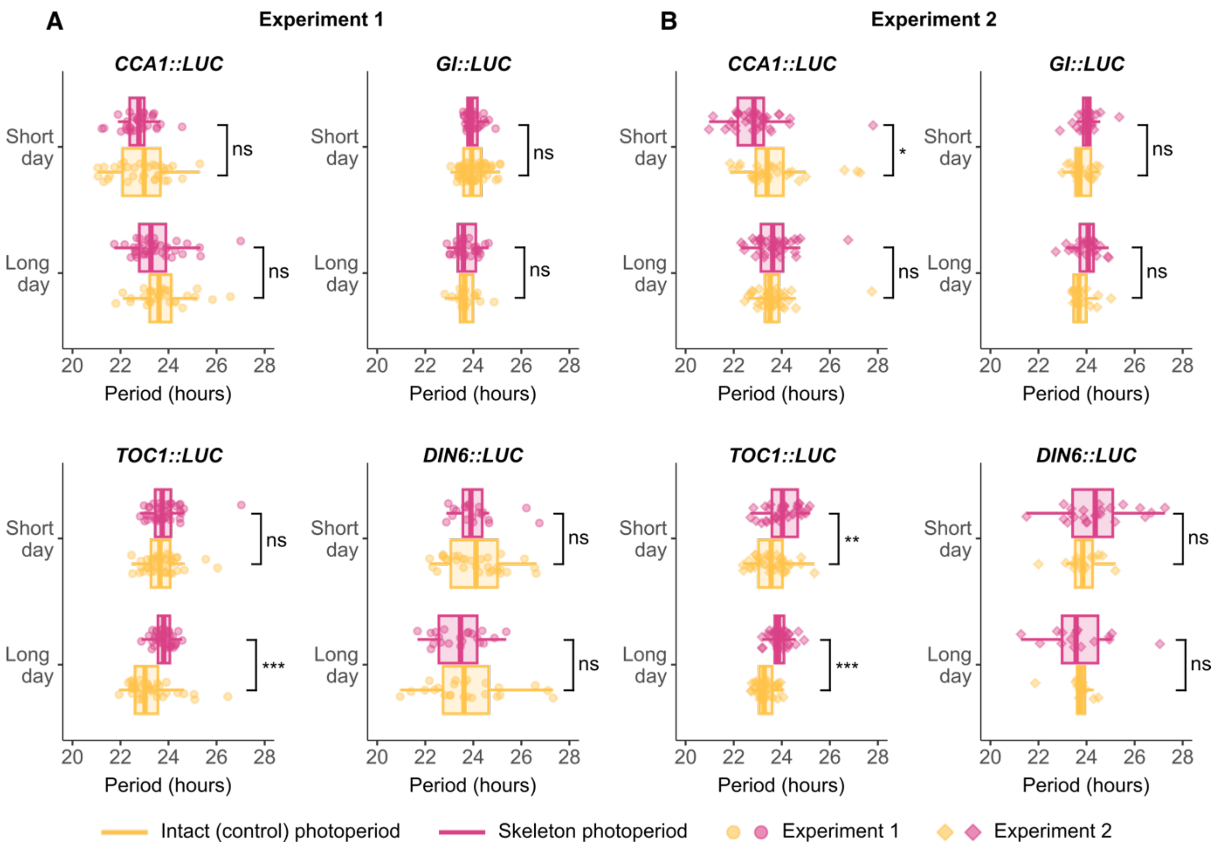


**Figure S6: Free-running period is unaltered by prior entrainment to skeleton photoperiods.** Free-running period of three circadian oscillator reporters (*CCA1::LUC*, *GI::LUC*, *TOC1::LUC*) and a downstream clock output gene (*DIN6::LUC*), determined using FFT-NLLS. Data derive from two independent experimental replicates; Experiment 1 **(A)** and Experiment 2 **(B)**. Period estimation used data collected between 24 - 72 h of constant light, following entrainment to short- or long-day conditions of intact or skeleton photoperiods. Statistical comparisons of mean period length between intact and skeleton photoperiods were conducted using unpaired two-tailed Student's t-tests: ns, not-significant; *, $p < 0.05$; **, $p < 0.01$; ***, $p < 0.001$ ($n$ = 10-61 individual seedlings).

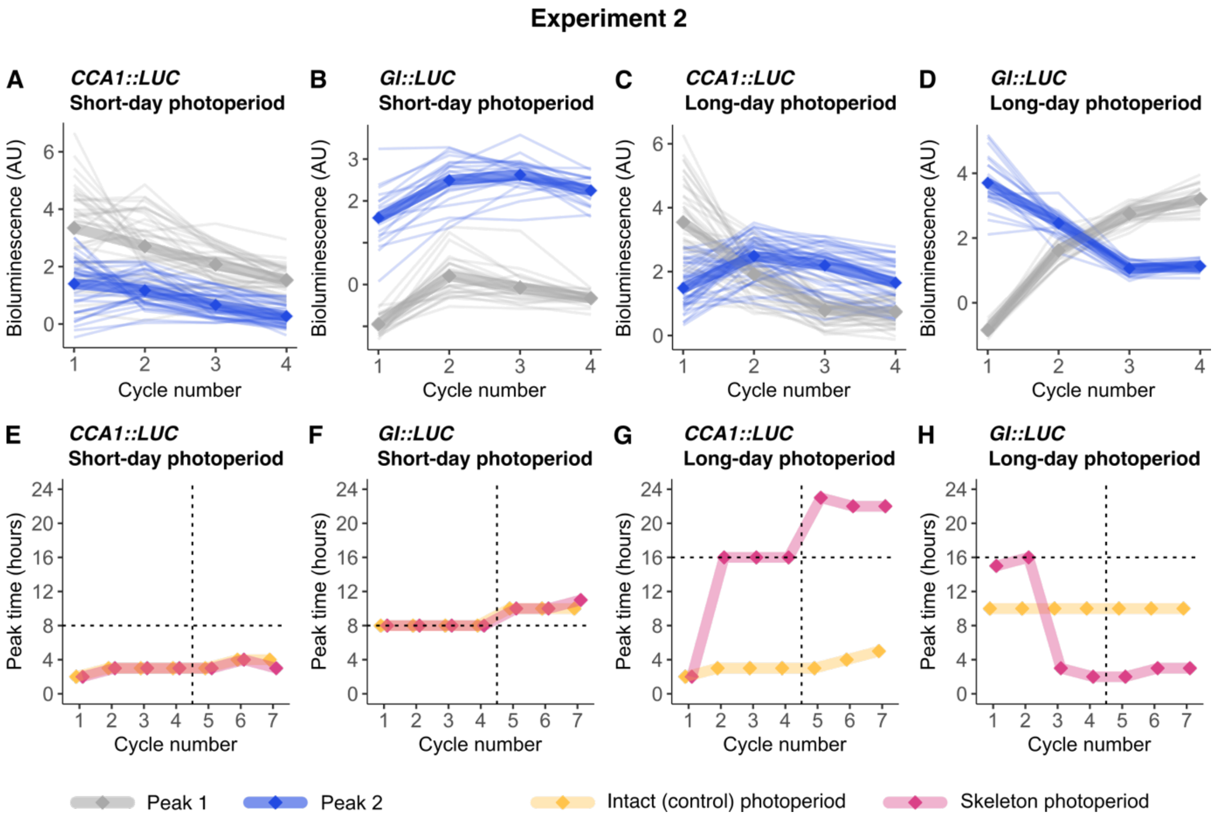


**Figure S7: Entrainment to long-day skeleton photoperiods switches circadian phase between the two light pulses. (A-D)** Peak bioluminescence measurements of *CCA1::LUC* and *GI::LUC* associated with dawn (peak 1; grey) dusk (peak 2; blue) light pulses, measured under both short- and long-day skeleton photoperiods**.** Each line represents an individual seedling ($n$ = 17-41) with the mean overlayed on top. **(E-H)** Examination of how these dynamics are reflected during entrainment and free-run under both intact and skeleton photoperiods. Data show timing of maximum daily bioluminescence of *CCA1::LUC* and *GI::LUC* across each 24 h window during both entrainment and subsequent free-run in constant light. Daily maxima may be due to either peak 1 or peak 2. Vertical dashed lines indicate the transition from entrainment to free-run: cycles 1-4 were the entrainment period, and cycles 5-7 were free-running conditions of constant light. Horizontal dashed lines indicate the end of the light period for intact photoperiods, and end of the second light pulse for skeleton photoperiods. Peak time data are mean of replicate seedlings. Data derived from an independent replicate experiment (Experiment 2).

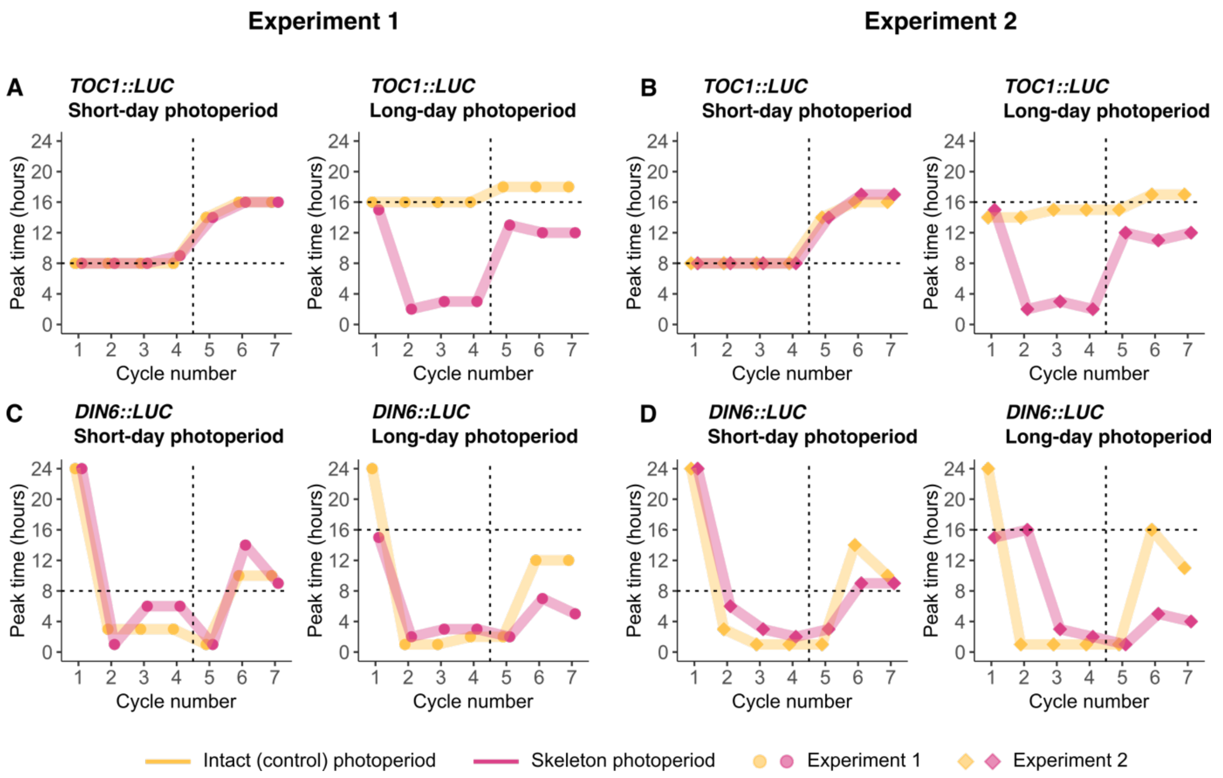


**Figure S8: Peak times of additional bioluminescence reporters under intact and skeleton photoperiods.** Timing of maximum daily bioluminescence of circadian clock component *TOC1::LUC* **(A, B)** and downstream clock output *DIN6::LUC* **(C, D)** across each 24 h window during both entrainment and subsequent free-run in constant light. Vertical dashed lines indicate the transition from entrainment to free-run: cycles 1-4 were the entrainment period, and cycles 5-7 were free-running conditions of constant light. Horizontal dashed lines indicate the end of the light period for intact photoperiods, and end of the second light pulse for skeleton photoperiods. Data are mean of replicate seedlings, from Experiment 1 **(A, C)**, and Experiment 2 **(B, D)**.

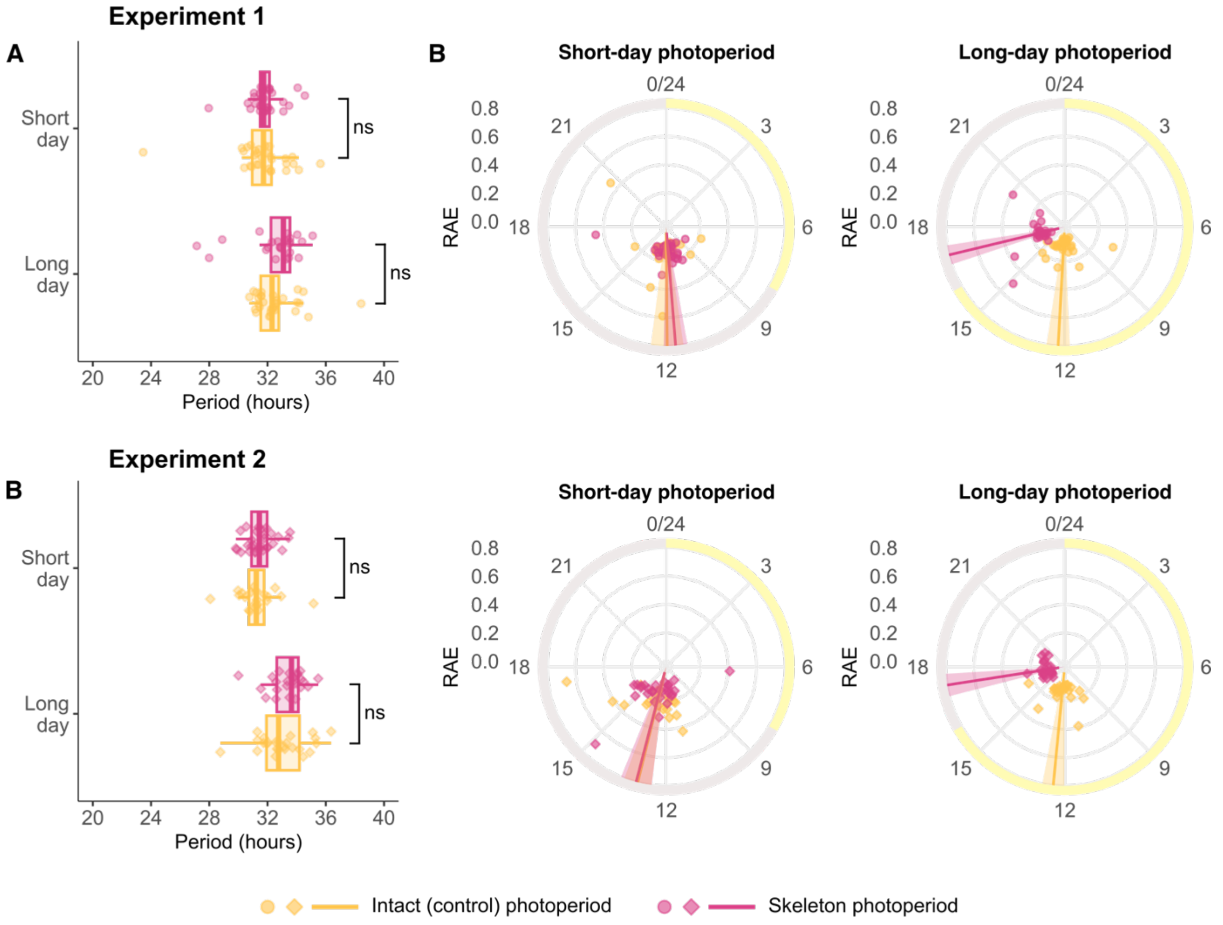


**Figure S9: Skeleton photoperiod-induced phase shifts of *GI::LUC* are also observed under free-running conditions of constant darkness.** Plants were entrained to short- or long-day conditions with intact or skeleton photoperiods and then transferred to free-running conditions of constant darkness. **(A)** Free-running period of *GI::LUC* under DD, with comparisons between entrainment conditions using unpaired two-tailed Student's t-tests (ns, not-significant). Period was estimated using FFT-NLLS. **(B)** Phase, derived from the free-run part of the data, shown as mean phase (line) ± angular deviation (ribbon; $n$ = 24-32 individual seedlings represented as points). Yellow shading around each phase plot indicates subjective day, and grey shading indicates subjective night. Radial axes indicate the relative amplitude error (RAE). Two independent experimental replicates are shown.

### Experiment 2

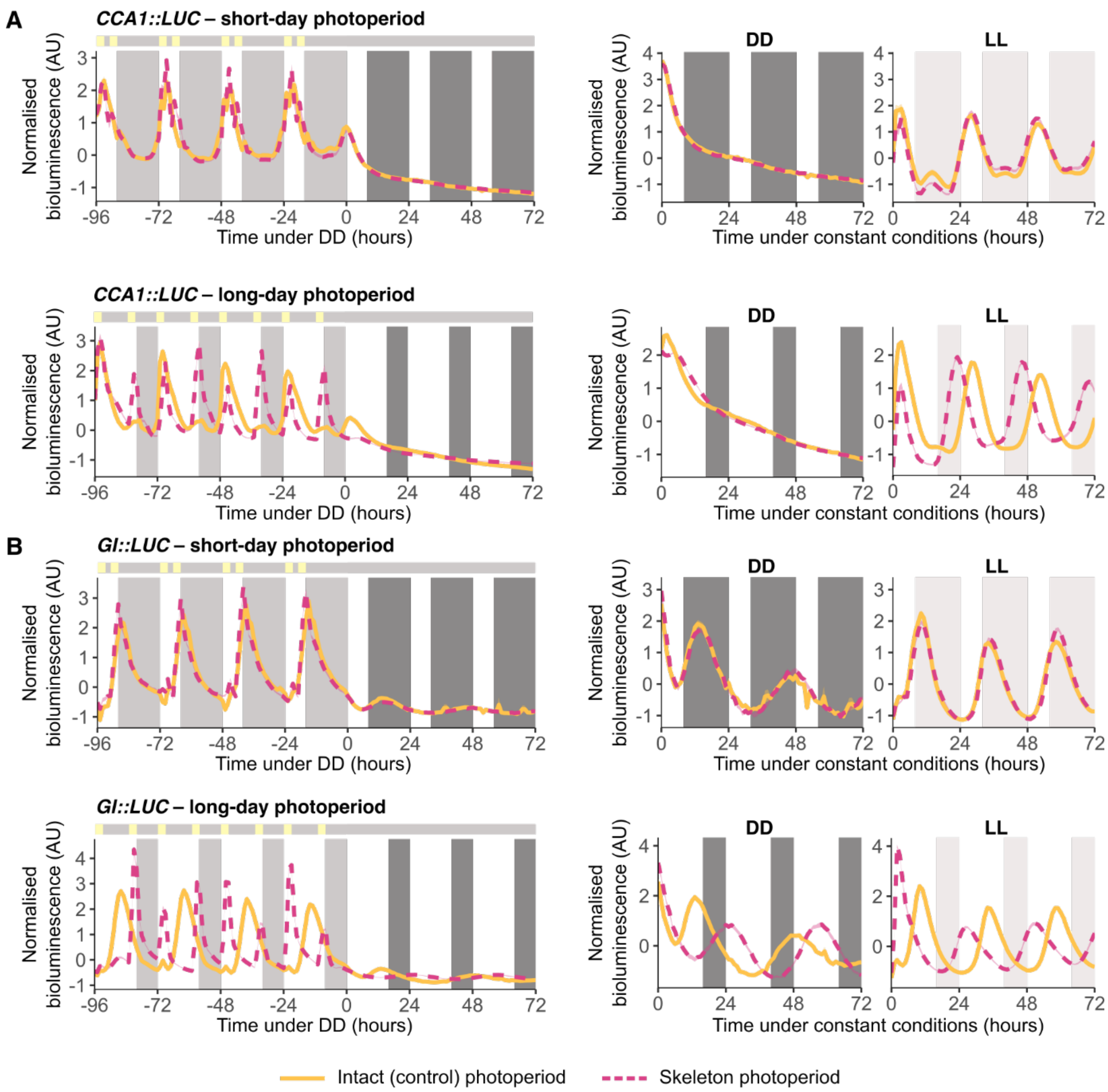


**Figure S10: The differing phases of clock reporters under long-day intact and skeleton photoperiods is confirmed under free-running conditions of constant darkness.** Normalised bioluminescence of *CCA1::LUC* **(A-D)** and *GI::LUC* **(E-H)** seedlings under short- **(A, B, E, F)** or long-day **(C, D, G, H)** conditions, using either intact (solid yellow) or skeleton (dashed pink) photoperiods. Following entrainment, plants were released into free-running conditions of constant darkness (DD) at ZT0. For panels **A**, **C**, **E,** and **G**, shading on the plot area indicates the light regime of the intact photoperiod, with light grey bars night during entrainment, and dark grey indicating subjective night under DD. The bar above each plot indicates the skeleton photoperiod regime, with yellow indicating lights on and grey indicating lights off. **(B**, **D**, **F**, **H)** Re-normalised data from the DD portion of **A, C, E, G**. Under DD *CCA1::LUC* in panels **B** and **D** was not rhythmic. LL sections of these panels are re-normalised data from Figure

S2, for comparison purposes. Data are mean (line) ± SEM (ribbon; *n* = 24-40 individual seedlings) from an independent experimental replicate.

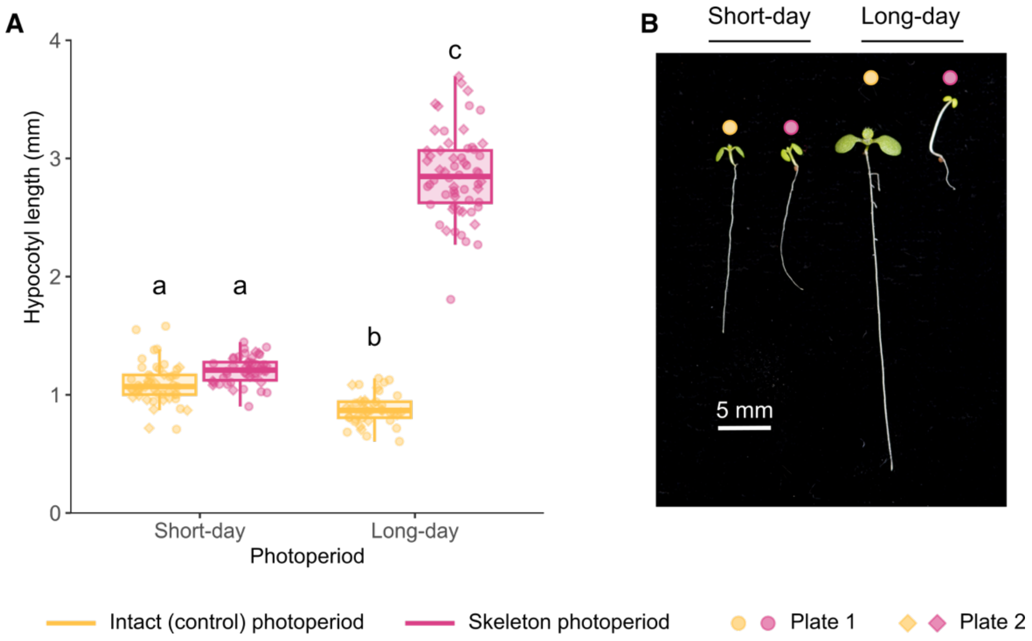


**Figure S11: Hypocotyl elongation under skeleton photoperiods. (A)** Hypocotyl lengths (mm) of Arabidopsis seedlings grown on two agar plates under constant light for 48 h followed by 7 days under either intact short-days (8L:16D), intact long-days (16L:8D), short-day skeleton (3L:2D:3L:16D), or long-day skeleton (3L:10D:3L:16D) photoperiods for 7 days. Statistical comparisons of hypocotyl length were conducted using a two-way ANOVA followed by Tukey's test for pairwise comparisons. Letters indicate statistical differences between groups ($p < 0.05$, $n$ = 19-33 individual seedlings). **(B)** Representative images of seedlings from each light treatment.
